# Are circulating levels of the myokine irisin linked to type 2 diabetes? A systematic review and meta-analysis

**DOI:** 10.1101/2024.06.29.601052

**Authors:** E. Aminov, P. Folan, A. Pisconti

## Abstract

**Background:** Type II diabetes (T2DM) is one of the most prevalent metabolic disorders, and its multisystemic health consequences are widely known. Due to skeletal muscle’s ability to sequester a vast amount of glucose, muscle function and exercise have become a subject of much research into strategies to prevent and treat T2DM. Myokines are bioactive molecules released by muscle during contraction and involved in several biological processes such as metabolism, inflammation and behavior. Irisin, a recently discovered myokine, has been implicated in a vast array of physiological roles, including the ability to induce fat beiging. Since beige and brown fat both serve important roles in metabolic regulation, irisin’s role in the context of T2DM is the subject of ongoing investigations.

**Methods:** We systematically reviewed articles indexed in PubMed, Scopus and Web of Science that were published between 2011 and 2024, and compared circulating irisin levels in patients affected by T2DM and healthy subjects. As part of our systematic review of the literature, we performed meta-analysis of the data across all included articles, as well as stratified by body mass index (BMI), country of origin and by average irisin concentration in the control group.

**Results:** We discovered great variability across the included studies in the average irisin levels detected, which spanned four orders of magnitude, hence the attempt at reducing variability by stratifying based on average levels in the control group. While the statistical power of our meta-analysis was decreased by the great variability in reported irisin concentrations, we nonetheless detected a consistent trend of decreased irisin concentration in T2DM patients compared with healthy controls, regardless of BMI, country of origin or average irisin concentration in the control group.

**Conclusion:** With almost 60 articles included, ours is the first extensive systematic review and meta-analysis of irisin in T2DM, yet a highly statistically significant association between circulating irisin levels and T2DM could not be established due to the great variability of the data across include articles. Nonetheless we noticed a trend that is independent of BMI, suggesting a direct relationship between T2DM and irisin that is likely not secondary to diabetic sarcopenia. While our work encourages further research into irisin’s potential role in T2DM pathogenesis, the reproducibility of irisin detection methods in biological samples should be determined and standardized protocols should be made available to the research and clinical communities.

## INTRODUCTION

Diabetes mellitus is a family of metabolic diseases marked by chronic hyperglycemia due to defects in insulin release and/or action. Both major types of diabetes mellitus, type 1 and 2 diabetes, often present similar symptoms at early stages including dysuria, polydipsia, weight loss, and blurred vision. Chronic hyperglycemia and progressed diabetes may accompany more severe symptoms such as diabetic retinopathy, blindness, susceptibility to infection, renal failure, increased risk of macrovascular complications, and increased risk of extremity amputation.

Type 1 diabetes (T1DM) is an autoimmune disorder most often marked by the chronic destruction of β-islets of Langerhans, endocrine cells specialized in the production and release of insulin. Although controversy exists on the mechanism by which β-cell destruction occurs, it is largely agreed upon that β-cell destruction and/or suppression is the reason for drastically lower blood insulin and drastically higher blood glucose levels in type 1 diabetics (DiMeglio et al., 2018). Despite the ongoing development of immune-altering drugs for T1DM, insulin analogs remain the most effective treatments currently in use (DiMeglio et al. 2018; Kahanovitz et al., 2017).

Type 2 diabetes (T2DM) has become one of the world’s most common diseases, with some measures indicating that approximately 8.3% of the global population meets the diagnostic criteria for T2DM (Kharroubi and Darwish 2015); additional estimates indicate that 10.1% of the population will suffer from T2DM by 2035 (Kharroubi and Darwish 2015). Although various subtypes of TD2M exist, the vast majority of type 2 diabetes cases result from insulin resistance, higher demand for insulin by the target tissue, drops in insulin secretion, and, possibly, eventual failure of β-cells (Halban et al. 2014).

The etiology of T2DM is complex and likely includes both genetic and lifestyle factors; significant research has been conducted to elucidate the effects of physical activity, diet, genetic predispositions, and cell signaling mechanisms on T2DM. Physical activity has been shown to cause acute and chronic increases in insulin sensitivity, lipid oxidation, hepatic and skeletal muscle glucose uptake, and β-cell mass in both healthy and diabetic subjects (Colberg et al., 2010; Kirwan et al., 2017). The most important risk factor identified for T2DM is obesity. Not only are the chances of developing T2DM positively associated with a high body mass index (BMI) but the severity of T2DM is also correlated with BMI; individuals with BMI >25 (obesity class 1) show the highest levels of HbA1c and mean fasting glucose (Nguyen et al., 2011). Family history of T2DM is another important risk factor; indeed genome-wide association studies (GWAS) have revealed over 100 loci that are associated with T2DM, many of which contain single-nucleotide polymorphisms, and T2DM concordance in twins is substantial (Kharroubi and Darwish 2015; Xue et al. 2018; Sabiha et al. 2021; Loh et al. 2022; Willemsen et al. 2015). From the signaling point of view, aPKC activation deficiencies are highly characteristic of Type 2 diabetics (Farese et al. 2005). Similarly, IRS-1 expression in cells from patients with non-insulin-dependent diabetes mellitus (NIDDM) is significantly reduced compared to healthy controls, while IRS-2 expression remains relatively constant between the two groups, signifying that IRS-1 expression may play an important role in the development of T2DM (Rondinone et al., 1997).

Management of T2DM is mostly reliant on lifestyle changes, but medications are available to those with severe conditions. Physical activity as well as proper dieting is associated with increased insulin sensitivity, prompting many healthcare providers to prescribe dieting and exercise routines to reverse T2DM (Hallberg et al. 2019).

Skeletal muscle’s metabolic functions, such as regulation of body temperature and glucose homeostasis, make it a prime target for T2DM treatments. As the largest glucose sink in the body, skeletal muscle accounts for the vast majority of glucose absorption from the bloodstream (∼85%; Bouzakri et al., 2005). Moreover, the rate of glycogenesis is limited by the speed at which skeletal muscle can absorb glucose (Cline et al., 1999).

Another important way in which skeletal muscle physiology is related to T2DM pathogenesis and progression is diabetic myopathy, a set of symptoms related to and caused by skeletal muscle deterioration as a consequence of diabetes. Reduced muscle strength and mass, and overall physical fitness are common manifestations of diabetic myopathy (Hernandez-Ochoa and Vanegas, 2015). Moreover, motor dysfunction and muscle atrophy are particularly exacerbated in distal portions of the legs and feet in T2DM patients that develop a neuropathy compared to non-neuropathic patients (Anderson et al., 1997). Lastly, increased levels of inflammatory signaling molecules in T2DM patients result in increased protein degradation, decreased protein synthesis, and eventually muscle atrophy (Giha et al., 2022).

Dysfunction of skeletal muscle glucose absorption mechanisms is perhaps the leading cause of insulin resistance in prediabetes and T2DM. It has been postulated that insulin resistance in skeletal muscle may be the initial cause of increased insulin secretion by β-cells (DeFronzo and Tripothy, 2009; Peterson and Shulman, 2002; Shou et al., 2020; Teng and Huang, 2019). A positive feedback cycle through which prediabetes can accelerate into T2DM with increased insulin resistance, leading to increased insulin release, can eventually terminate in the partial or total failure of β-cells (Cerf, 2013; Halban et al., 2014; Kasuga, 2006).

Glucose consumption is not the only mechanism through which skeletal muscle activity participates in metabolic regulation; many autocrine, paracrine, and endocrine signaling molecules are released into the interstitial fluid and circulation, which are active in the regulation of metabolic processes. These signaling molecules have been termed myokines (Lee and Jun, 2019; Severinson and Pederson, 2020). By comparing pre and post-exercise human plasma, it has become apparent that myokine release can be stimulated by acute exercise as well as long-term training regimens (Hoffman and Weigert, 2017; Jedrychowski et al., 2017; Ostrowski et al., 1998). IL-6, most commonly known for its role as a regulator of inflammation (Tanaka et al., 2014), is also a myokine with a wide range of metabolic effects (Rehman et al., 2017; Simpson et al., 1997; Ostrowski et al., 1998) and is increased in plasma post-acute exercise (Hoffman and Weigert, 2017). As one of the most studied myokines, IL-6 has many metabolic functions, some of which are directly related to the management of blood glucose levels and may play important roles in the prevention of T2DM. IL-6 stimulates glucose disposal, glucose oxidation, lipolysis, and fat oxidation (Carey et al., 2006; van Hall et al., 2003). In addition to IL-6, other myokines, such as such as myostatin, IL-15 and ANGPTL4, involved in glycemic control, body fat content, oxidative potential, and muscle mass have been identified (Hittel et al., 2010; Kersten et al., 2009; Riechman et al., 2004).

Irisin is a myokine derived from the cleavage of fibronectin type III domain-containing protein 5 (FNDC5) in skeletal muscle following acute exercise (Boström et al. 2012; Arhire et al. 2019). In mouse models, acute exercise and various training regimens result in increased levels of irisin; the same phenomenon can be observed in humans but with less consistency (Boström et al. 2012; Pekkala et al. 2013; Wrann et al. 2013). Importantly, irisin promotes skeletal muscle hypertrophy by promoting muscle stem cell (satellite cell) activation and directly enhancing protein synthesis (Reza eta l., 2017). Moreover, irisin rescues skeletal muscle atrophy in a model of muscle denervation (Reza et al., 2017), further stressing its importance as a myokine with both paracrine and autocrine activity.

Irisin has many important metabolic functions (Fig. 1), for example it is a key hormone in the process of fat “beiging” through which white adipose tissue (WAT), functioning primarily in the storage of excess triglycerides, turns into beige fat, which instead is instrumental to the regulation of the metabolic rate through the uncoupling of mitochondrial respiration and thermogenesis (Harms and Seal 2013). The word “beiging” is meant to describe the way in which white fat cells switch to a mechanism of thermoregulation that is similar to that used by brown fat cells (Wu et al. 2013). Additionally, autocrine irisin signaling in skeletal muscle is associated with increased glucose uptake, gluconeogenesis, lipid metabolism, and lipid uptake (Arias-Loste et al. 2014; Liu et al. 2017).

**Figure 1.**
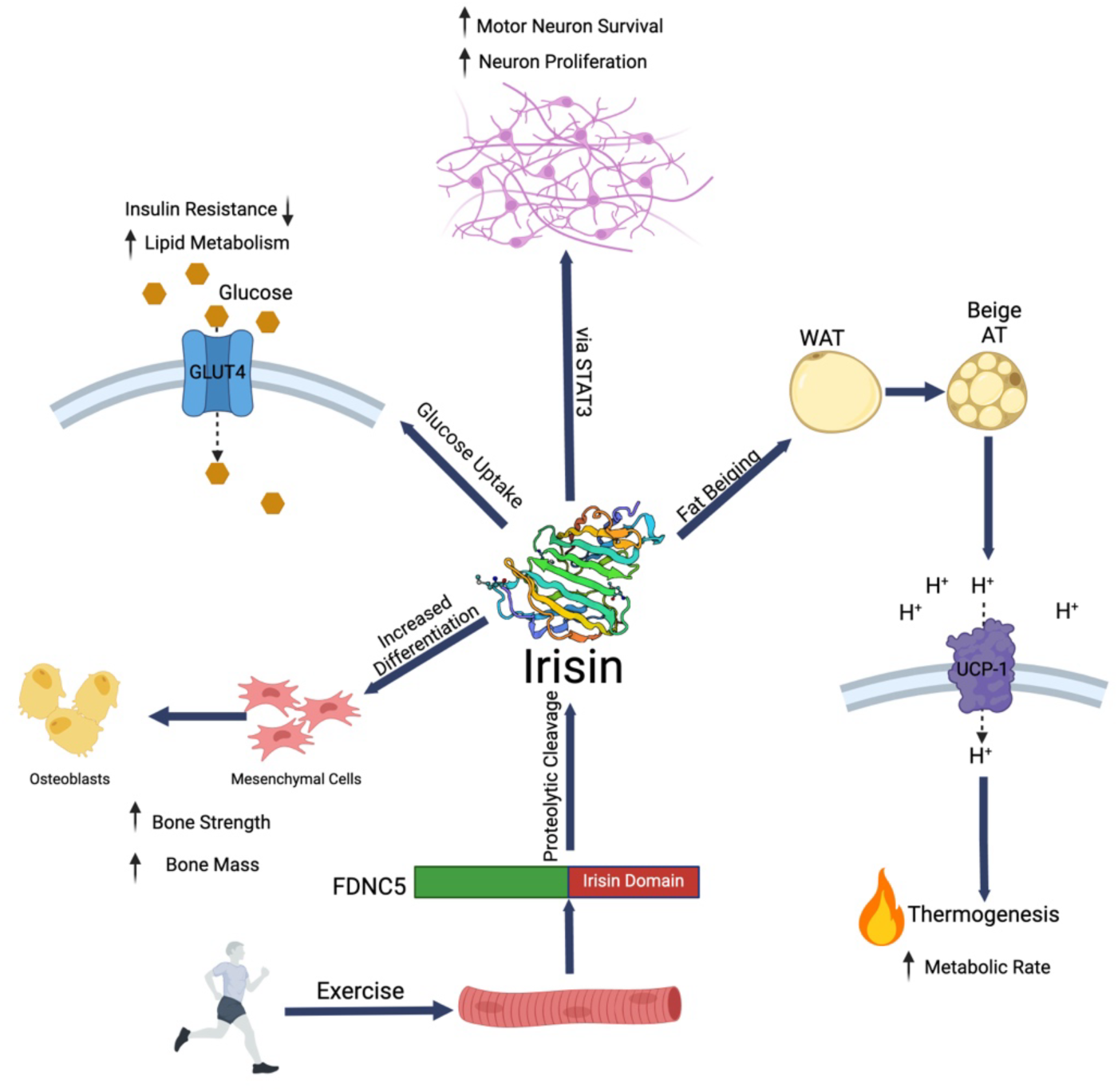
Irisin is a myokine produced by cleavage of FDNC5 during acute exercise and is involved in a plethora of biological processes.

Irisin has also been associated with many functions other than those directly related to skeletal muscle and adipose tissue (Fig. 1). *In vitro* and *in vivo* studies have uncovered irisin’s relationship to bone, the nervous system, and chronic inflammation (Korta et al. 2019). *In vitro,* irisin improves osteoblast differentiation and increases the levels of osteoblastic transcription regulators (Qiao et al. 2016). In mouse hippocampal neurons, irisin induces cell proliferation via STAT3 (Moon et al. 2013), a transcriptional regulator implicated in various pathways associated with nervous system development, increased motor neuron survival and the differentiation of glial cells (Dziennis 2008). Moreover, mouse ischemia models have shown that irisin has significant neuroprotective effects and is associated with decreased brain edema, neurological deficits, and brain infarct volume post-ischemia (Li et al., 2017). Irisin also has anti-inflammatory properties (Fig. 1), likely acting by downregulating the expression of pro-inflammatory cytokines and Toll-like receptor 4 (TLR4), as well as via reducing the production of hydrogen peroxide and reactive oxygen species (Korta et al. 2019).

Due to T2DM’s metabolic nature and association with a state of chronic inflammation, it is reasonable to postulate that irisin may play a role in T2DM pathogenesis. Studies comparing irisin levels in patients affected by T2DM and healthy subjects found that patients with T2DM had significantly lower circulating levels of irisin (Du et al., 2016). Although findings of lower irisin levels in T2DM patients are consistent, the role of irisin in T2DM pathogenesis and treatment remains elusive. For example, it is unknown if lower reported irisin levels in T2DM patients are due to insulin resistance in skeletal muscle and subsequent diabetes-induced muscle atrophy, or if lower irisin levels may act as a causal factor in the development of T2DM. Most importantly, potential confounding factors have not been systematically investigated.

## OBJECTIVE

The objective of this article is to systematically review the current literature on circulating irisin levels in patients affected by T2DM and use meta-analysis to investigate the relevance of potential confounding factors such as BMI and country of origin.

## METHODS

### Literature Search

The PubMed, Web of Science (WOS), and Scopus databases (2011-2024) were searched for relevant studies. The initial scoping search query was (irisin[Title/Abstract]) AND (Diabetes[Title/Abstract]). Results from this scoping search were refined to exclude narrative reviews, papers published before January 1st, 2011 and after February 12th, 2024, and studies on forms of diabetes other than T2DM. The final search query was as follows for PubMed: (((“Irisin”[Title/Abstract] AND “Diabetes”[Title/Abstract]) NOT “review”[Filter]) NOT “type 1”[Title]) NOT “Gestational”[Title]”. Search queries of identical criteria were administered for the Web of Science and Scopus databases. Before screening, duplicate articles were removed from the results by comparing titles of articles retrieved from each database (Figure 2). After results were obtained, abstracts and titles were screened, and inclusion/exclusion criteria were applied. All references of included papers were scanned for additional studies. Each article’s eligibility for inclusion was assessed after applying inclusion/exclusion criteria by reading the full texts. Scoping searches were completed on February 12th, 2024. All peer-reviewed English language material comparing irisin levels in adult humans with T2DM to healthy adult controls with normal glucose tolerances were included in the metanalysis, including cross-sectional studies, case-control studies, longitudinal studies, and clinical trials. Studies were excluded if they were not peer-reviewed, lacked either a T2DM or control group, included prediabetes/metabolic syndrome subjects in either experimental group, combined T2DM groups with gestational or Type 1 diabetes subjects, or included children. Obese controls were included if they were normal glucose tolerant. Systematic and narrative reviews were excluded from metanalysis, but original studies analyzed in other systematic reviews were included.

**Figure 2:**
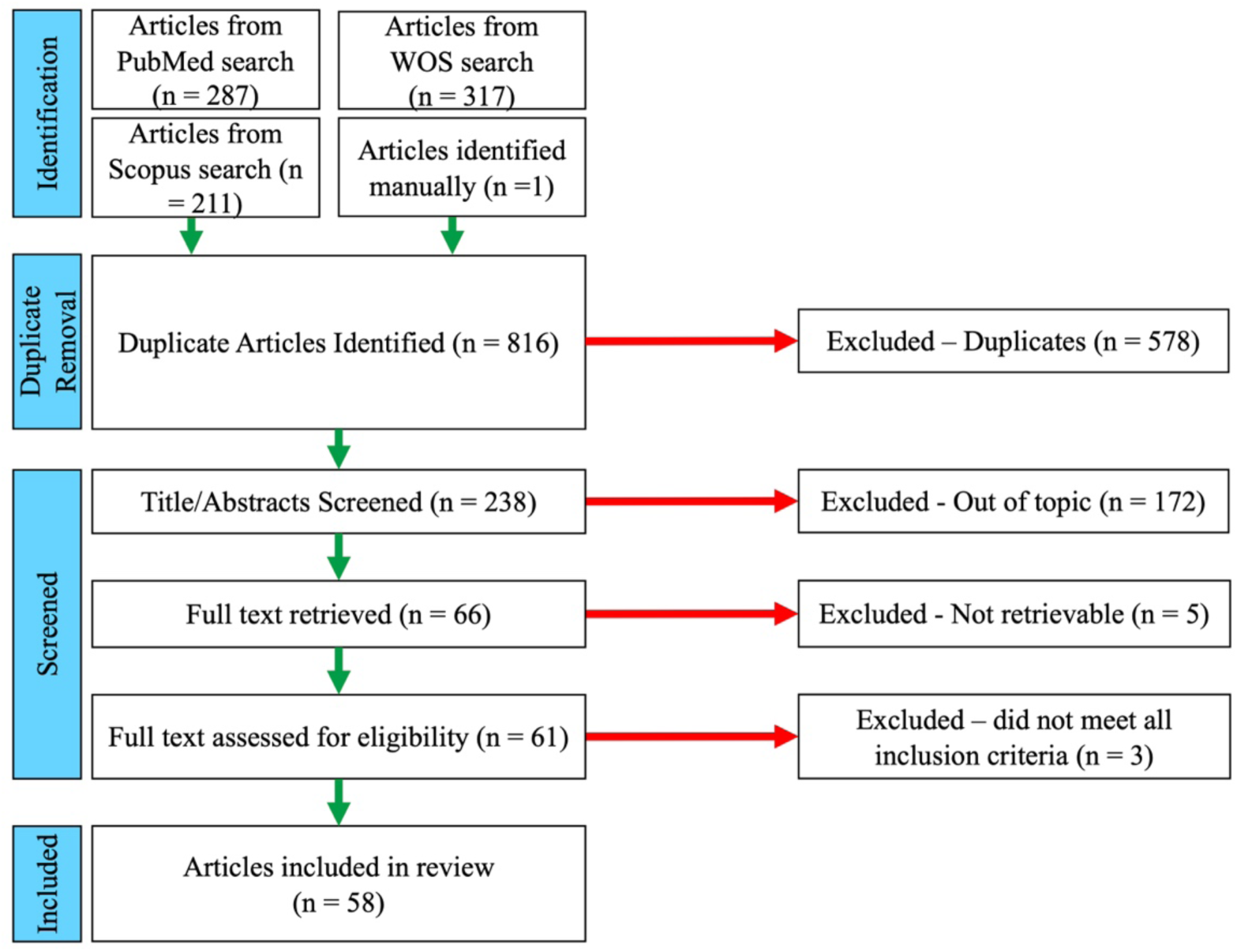
PRISMA chart of article selection.

Searching the PubMed, WOS, and Scopus databases using the aforementioned queries yielded 287, 317, and 211 articles, respectively. An additional article was identified manually in the references of a systematic review by Song et al. 2021. Duplicate identification resulted in the exclusion of 578 articles. After initial screening and application of inclusion and exclusion criteria, 172 articles were excluded. 66 articles were sought for retrieval and 61 were assessed for eligibility. One article was excluded after eligibility assessment due to HbA1c averages of both diabetics and healthy controls exceeding well past abnormal limits (54.6% for controls and 66.02% for diabetics; Choi et al. 2013, while the average HbA1c range for a healthy individual is less than 5.7%, CDC source). Two articles were excluded due to the inclusion of adolescent participants, and an additional four articles were inaccessible. A total of 58 articles were included in the meta-analysis comparison of irisin levels (Fig. 2).

### Data Collection

Data were collected from included studies, and conflicting data units and formats were resolved. If skewed data were presented as median and interquartile range (IQR), mean and standard deviation were estimated (Wan 2014). Where necessary, continuous data from different groups were combined using the group combination equation from Cochrane (Higgins et al. 2022). After data collection was performed on all included studies, irisin levels and sample sizes for both normal glucose tolerant and T2DM groups were transferred into Cochrane’s Review Manager, RevMan Version 5.4.1, for metanalysis.

### Meta-Analysis

Irisin levels of T2DM patients were compared against irisin levels of healthy controls in a meta-analysis across all included studies. The standardized mean difference was used to generate forest plots with a fixed effect analysis method (Higgins et al. 2022). 95% confidence intervals were calculated by RevMan Version 5.4.1 to estimate the true effect size of T2DM on irisin levels. Heterogeneity of the outcomes was assessed by calculating the inconsistency index, *I^2^*. The heterogeneity of the included studies was high due to different articles reporting greatly divergent (in the order of several orders of magnitude) average levels of irisin in both non-diabetic and T2DM cohorts. We were not able to establish a significant correlation between the average irisin levels reported and any of the methods used in each article which could have introduced analytical biases in irisin quantification. For this purpose, we assessed the correlation between average irisin levels in the non-diabetic cohort and:

a. Specific Enzyme-linked immunosorbent assay (ELISA) kit or antibody used to quantify irisin.
b. Whether plasma was used or serum.
c. Biological sex of participant.
d. Body mass index.
e. Triglyceride levels.

None of these attempts yielded significant correlation, largely due to the great variability within the irisin levels dataset. Thus, we supplemented the meta-analysis of all included studies with meta-analyses of subgroups stratified by:

i) Absolute irisin levels. Stratification of included studies based on maximum irisin levels observed in the non-diabetic cohort was conducted as follows, leading to five subgroups:
  1. max irisin levels in non-diabetics <1.00 ng/mL (n = 5 studies; 847 subjects)
  2. max irisin levels in non-diabetics = 1.00 – 50.00 ng/mL (n = 19 studies; 3139 subjects)
  3. max irisin levels in non-diabetics = 50.01 – 100.00 (n = 4 studies; 579 subjects)
  4. max irisin levels in non-diabetics = 100.01 – 500.00 ng/mL (n = 16 studies; 1876 subjects)
  5. max irisin levels in non-diabetics > 500.01 ng/mL (n = 14 studies; 1549 subjects) Additional subgroup analysis was performed to control for potential confounding factors through stratification of studies by:
ii) Country in which the study was conducted, due to known differences in the prevalence of diabetes in different countries (Lin et al. 2022). If greater than five studies were conducted in a country, a separate subgroup was created. Otherwise, studies were included in the “other” subgroup. Subgrouping by country yielded three subgroups:
  1. China (23 studies; 3879 subjects)
  2. Egypt (9 studies; 1384 subjects)
  3. Other (26 studies, 2852 subjects).
iii) BMI class, due to the association between BMI class and T2DM (Bays et al. 2007), Subgrouping based on the BMI classification of the control group was conducted as follows according to CDC definition of BMI classes:
  1. Normal: BMI = 18.5 – 24.9 (23 studies, 3107 subjects)
  2. Overweight: BMI = 25.0 – 29.9 (20 studies 3046 subjects)
  3. Obese: BMI ≥ 30 (6 studies, 852 subjects)

Lastly, subgrouping was also performed to compare studies reporting continuous data with studies reporting parametric data that had been converted to mean ± standard deviation, to determine if data conversion had a significant impact on calculated effect sizes. Conversion of parametric data to continuous data was achieved using methods established by Wan 2014. The continuous data subgroup contained 47 studies with a total of 6569 subjects. The parametric-converted subgroup contained 11 studies with a total of 1546 subjects.

## RESULTS

Table 1 summarizes all the included studies and the irisin levels detected in the two populations studied – T2DM patients and healthy control (non-diabetic) participants – as well as other relevant data including: country where the study was conducted, the diagnostic criteria followed in defining the T2DM population, the ELISA kit used to assess circulating levels of irisin, sample size for both cohorts (case and control), Hba1c levels, and BMI.

**Table 1.** Overview of included studies sorted by reported irisin concentrations. Data are presented as mean ± standard deviation. Breaks in the table indicate which absolute irisin stratification group each study belonged to in the meta-analysis.

| First Author/<br>Year (irisin<br>subgroup) | Country | Diagnostic<br>criteria | Irisin<br>ELISA Kit | N (case/<br>control) | % HbA1c<br>(case/ctrl) | Fasting<br>glucose<br>(mmol/L) | Irisin (ng/mL)<br>(case/control) | BMI<br>(case/control) |
| --- | --- | --- | --- | --- | --- | --- | --- | --- |
| Akgul<br>Balaban 2019<br>(5) | Turkey | N. D. | Hangzhou<br>Eastbioph. | 41/49 | 8.5 ± 2.7/<br>5.2 ± 0.6 | 9.9±0.2/<br>4.8±0.5 | 2790 ± 830/<br>3340 ± 970 | 29.8 ± 6/<br>26.5 ± 3.5 |
| Al-Daghri<br>2015 (5) | Saudi<br>Arabia | N. D. | Phoenix<br>Pharm. | 83/81 | N. D. | 12.7±3.6<br>/4.8±1.1 | 890 ± 30/<br>590 ± 30 | 29.4 ± 4.7/<br>29.6 ± 4.3 |
| Ali 2020 (5) | Saudi<br>Arabia | ADA | BioVendor | 92/46 | 8.1 ± 1.7/<br>5.3 ± 0.3 | 9.8±3.1/<br>5.1±0.6 | 540 ± 140/<br>650 ± 310 | 29.47 ± 5.44/<br>30.22 ± 5.29 |
| AlKhairi<br>2019 (5) | Kuwait | ADA (2019) | Phoenix<br>Pharm. | 104/1<br>23 | 7.7 ± 0.2/<br>5.6 ± 0.1 | 8.2±0.3/<br>4.8±0.5 | 623 ± 223/<br>508 ± 171 | 31.62 ± 0.42/<br>28.86 ± 0.48 |
| Duran 2015<br>(5) | Turkey | ADA (2006) | Adipogen | 50/52 | N. D. | 7.7±2.4/<br>5.0±0.4 | 2830 ± 520/<br>3160 ± 300 | 32.38 ± 4.4/<br>30.54 ± 5.9 |
| Ebert 2016<br>(5) | Germany | ADA (2009) | Adipogen | 29/33 | N.D. | N.D. | 813.3 ±<br>179.3/ 986.6<br>± 348.7 | N.D. |
| Feng 2015 (5) | China | WHO | Phoenix<br>Pharm. | 25/40 | 6.9 ± 0.7/<br>5.4 ± 0.3 | 7.43±1.0<br>1/5.06±0<br>.50 | 45.15±10.48<br>/35.38±9.97 | 26.27±4.6/<br>24.63±3.71 |
| Guo 2020 | China | WHO<br>(1999) | Sino Best<br>Biol. Tech. | 71/71 | N.D. | 7.8±1.74<br>/5.2±0.4 | 640.7±100.1<br>/780.2±29.3 | 22.17±2.4/<br>21.1±2.08 |
| Huh 2022 (5) | S. Korea | WHO | Phoenix<br>Pharm. | 85/85 | 6.0 ± 0.3/<br>5.5 ± 0.3 | 5.6±0.5/<br>5.0±0.5 | 2282±1291/<br>1585±830 | 26.27±3.12/<br>24.52±3.29 |
| Komosinska<br>2020 (5) | Poland | Polish Diab.<br>Association | BioVendor | 40/20 | 6.3 ± 0.6/<br>5.2 ± 0.3 | 7.4±0.6/<br>5.0±0.3 | 8950 ±<br>5198/ 4813<br>± 2290 | 32.76 ± 4.83/<br>29.59 ± 6.22 |
| Mathia 2023<br>(5) | Brazil | HbA1c >7%<br>for at least 5<br>years | Phoenix<br>Pharm. | 16/14 | 7.1 ± 1.6/<br>5.6 ± 0.5 | 7.9±2.2/<br>5.1±0.1 | 498300±763<br>00/448600±<br>198300 | 27.6 ± 4.3/<br>26.4 ± 4.5 |
| Moreno-Naverrete<br>2013 (5) | Spain | ADA | Aviscera<br>Biosci. | 7/69 | N. D. | 6.45±0.6<br>1/5.31±.<br>53 | 1508 ± 222/<br>1849 ± 508 | 29.58 ± 3.4/<br>27.61 ± 3.8 |
| Shanaki 2017<br>(5) | Iran | ADA | BioVendor | 41/40 | N. D. | 8.3±2.9/<br>5.1±0.5 | 2200 ±<br>1540/ 4390<br>± 2800 | 26.91 ± 0.65/<br>24.04 ± 0.73 |
| Xuan 2020<br>(5) | China | ADA (2011) | Sino Best<br>Biol. Tech. | 71/71 | N. D. | N.D. | 703 ± 242/<br>800 ± 276 | 21.2 ± 2.3/<br>23.7 ± 3.5 |
| Alis 2014 (4) | Spain | ADA (2014) | Phoenix<br>Pharm. | 69/75 | 7.1 ± 1.0/<br>5.5 ± 0.3 | 8.0±2.5/<br>5.2±0.5 | 263 ± 38/<br>279 ± 58 | 30.6 ± 5.1/<br>33.0 ± 8.8 |
| El-Lebedy<br>2018 (4) | Egypt | ADA | MyBio-Source | 135/7<br>0 | 6.7 ± 1.2/<br>5.4 ± 0.5 | 8.8±3.4/<br>4.4±0.5 | 85.9 ± 68/<br>127 ± 72 | 24.05 ± 3.17/<br>22.8 ± 2.2 |
| Garcia-Fontana 2015<br>(4) | Spain | ADA (2005) | Phoenix<br>Pharm. | 73/55 | 8.0 ± 1.9/<br>4.9 ± 0.4 | 9.9±3.3/<br>5.0±0.6 | 462 ± 91/<br>429 ± 73 | 31 ± 6/<br>29 ± 6 |
| He 2015 (4) | China | N. D. | Phoenix<br>Pharm. | 72/28 | 8.2 ± 2/ 5.8<br>± 0.4 | 8.0±3.0/<br>5.3±0.6 | 180 ± 24/<br>427 ± 95 | N. D. |
| Jasim 2022<br>(4) | Iraq | N.D. | N.D. | 90/25 | 2.9±2.2/<br>2.5±2.2 | N.D. | 27.07±9.6/<br>151±5.9 | N.D |
| Khajebishak<br>2023 (4) | Iran | N.D. | Zellbio<br>GmbH | 32/30 | N.D. | 7.5±2.1/<br>4.8±0.3 | 218.7±50.5/<br>266.7±69.89 | 34.35±4.07/<br>24.33±3.7 |
| Khidr 2017<br>(4) | Egypt | ADA (2014) | BioVision | 100/5<br>0 | 10 ± 0.6/ 5.2<br>± 0.1 | 12.2±0.4<br>/5.1±0.1 | 203 ± 58/<br>315 ± 7 | 24.2 ± 0.24/<br>23.9 ± 0.1 |
| Liu 2013 (4) | Singapore | ADA (2006) | USCN Life Science | 96/60 | 8.3 ± 1.9/N.D. | N.D/5.3 ±0.8 | 204 ± 72/257 ± 24 | 27.8 ± 4.6/25.0 ± 4.8 |
| Manjun 2022 (4) | Iraq | N.D. | Elabscience | 40/40 | N.D. | N.D. | 183.3±104.3/349±108.6 | N.D. |
| Montazerifar 2022 (4) | Iran | ADA | Bioassay Tech. Lab | 40/40 | 7.75±1.7/1.80±0.14 | 9.9±2.6/4.7±0.6 | 294.6±36/152.5±13.4 |  |
| Park 2020 (4) | South Korea | ADA | Phoenix Pharm. | 71/19 | 7.3 ± 1.6/5.3 ± 0.3 | 7.8±2.3/5.2±0.3 | 128 ± 38/176 ± 46 | 24.4 ± 2.9/24.1 ± 2.7 |
| Shoukry 2016 (4) | Egypt | ADA (2014) | BioVision | 150/150 | 9.0 ± 1.0/5.2 ± 0.3 | 9.20±0.7/4.9±0.3 | 273.1 ± 21.2/ 377.8 ± 27.2 | 39.15 ± 3.58/32.36 ± 6.87 |
| Tarboush 2021 (4) | Jordan | N. D. | Phoenix Pharm. | 25/34 | 8.0 ± 0.4/6.0 ± 0.1 | N.D. | 138 ± 8/294 ± 13 | 29.0 ± 1.0/26.0 ± 0.5 |
| Xie 2015 (4) | China | ADA | Phoenix Pharm. | 50/50 | 6.9 ± 1.9/5.4 ± 0.3 | 7.5±2.3/5.2±0.4 | 269.8±143.5/258.7±102 | 24.71±1.75/24.45±2.06 |
| Yang 2014 (4) | China | WHO | Phoenix Pharm. | 60/62 | 7.0 ± 1.8/5.5 ± 0.3 | 7.8±2.5/5.13±0.47 | 298.9 ± 141/277.6 ± 131.1 | 25.8 ± 2.99/24.07 ± 3.21 |
| Zhang 2016 (4) | China | ADA | Phoenix Pharm. | 50/50 | N. D. | 9.87±3.54/5.13±0.39 | 231 ± 44/253 ± 26 | 23.85 ± 2.33/23.72 ± 2.32 |
| Aly 2020 (3) | Egypt | FBG ± 140 mg/dL | DRG | 120/60 | N. D. | 15.4±0.1/4.9±0.0 | 86.99 ± 27.7/ 60.22 ± 12.94 | N. D. |
| Liu 2016 (3) | China | ADA (2013) | Aviscera Biosci. | 54/38 | 9.8 ± 2.1/5.8 ± 0.6 | 9.63±3.91/5.43±0.47 | 40.9 ± 18.7/62.96 ± 34.45 | 32.15 ± 5.07/32.29 ± 6.15 |
| Mostafa 2021 (3) | Egypt | WHO | Sunred Biological Tech. | 60/60 | 8.1 ± 1.5/5.5 ± 0.6 | 9.8±1.0/4.9±0.5 | 66.6 ± 16.2/88.4 ± 21.5 | 25.79 ± 2.52/25.44 ± 2.53 |
| Sahin-Efe 2018 (3) | USA | N. D. | Aviscera Biosci. | 47/140 | 6.0 ± 0.4/5.4 ± 0.4* | 7.8±0.2/5.5±0.1 | 79.64 ± 4.03/ 84.31 ± 2.34 | 28.9 ± 0.5/27.4 ± 0.3 |
| Ahmed 2023 (2) | Egypt | N.D. | N.D. | 65/25 | 9.1±1.6/5.1±0.67 | 10.3±2.4/4.8±0.6 | 3.34±1.9/3.36±1.3 | 25.5 ±3 .7/22.6 ± 2 .4 |
| Al-Aridhi 2022 (2) | Iraq | N.D. | Biosource | 88/25 | 8.4±2.1/5.4±0.53 | 12.0±4.5/5.3±0.4 | 7.2±3.2/30.15±16.03 |  |
| Berezin 2022 (2) | Ukraine | ADA (2014) | Elabscience | 183/25 | 6.6 ± 0.6/4.2 ± 0.95 | 5.6±1.3/4.2±0.7 | 6.29±3.54/15.1±0.7 | 25.5±3.7/22.6±2.4 |
| Hashim 2022 (2) | Iraq | N.D. | Sino Best Biol. Tech. | 60/60 | 8.3 ± 1.8/5.8 ± 0.51 | N.D. | 14.45±3.0/4.1±0.31 | N.D. |
| Li 2016 (2) | China | N. D. | Phoenix Pharm. | 20/13 | 11 ± 2/ 5.5 ± 0.4 | 9.9±3.3/5.2±0.6 | 15.4 ± 6.6/20.4 ± 6.8 | 25.9 ± 4/23.1 ± 2.4 |
| Li 2017 (2) | China | WHO (1999) | Aviscera Biosci. | 362/100 | 7.7 ± 1.2/5.3 ± 0.5 | N.D. | 16.24 ± 5.16/ 24.35 ± 2.76 | 25.7 ± 3/23.9 ± 3.4 |
| Li 2024 (2) | China | WHO (2019) | N.D. | 77/18 | 8.7 ± 1.8/5.5 ± 0.3 | 9.3±2.8/5.5±0.3 | 30.03±44.5/117.4±65.97 | 26.17±3.48/23.41±0.63 |
| Tang 2015 (2) | China | WHO (1999) | Phoenix Pharm. | 72/68 | N. D. | 7.14±1.05/5.19±0.56 | 10.6 ± 14.3/10.62 ± 13.52 | 25.21 ± 2.93/25.46 ± 2.98 |
| Nakamura 2017 (2) | Japan | JDS | Phoenix Pharm. | 18/12 | 7.4 ± 0.8/5.3 ± 0.2 | 6.9±0.7/4.84±0.4 | 12.2 ± 4.8/10.3 ± 1.2 | 26.5 ± 4.0/22.0 ± 3.1 |
| Wang 2015 (2) | China | ADA (2012) | Aviscera Biosci. | 100/100 | 9.9 ± 1.8/4.5 ± 1.6 | 8.9±1.5/4.8±1.1 | 14.12 ± 3.93/ 28.98 ± 2.56 | 27.8 ± 3.5/22.7 ± 5.6 |
| Wang 2016A (2) | China | WHO (1999) | Phoenix Pharm. | 100/20 | 8.3 ± 1.9/5.3 ± 0.3 | N.D. | 3.3 ± 1.07/4.78 ± 0.77 | 26.5 ± 4.43/26.21 ± 2.15 |
| Wang 2016B (2) | China | WHO (1999) | Wuhan Eiaab Science | 120/20 | 7.8 ± 0.3/5.3 ± 0.1 | 7.8±0.3/5.3±0.1 | 3.4 ± 0.1/4.7 ± 0.1 | 26.3 ± 0.4/26.0 ± 0.5 |
| Wang 2021 (2) | China | ADA (2014) | Phoenix Pharm. | 32/20 | 8.0 ± 0.8/5.6 ± 0.5 | 8.51±1.71/4.97±0.59 | 10.03 ± 2.06/ 13.03 ± 3.1 | 26.16 ± 3.44/25.24 ± 3.59 |
| Wang 2022 (2) | China | N. D. | Westang | 66/82 | 8.4 ± 3.6/<br>5.4 ± 0.2 | 8.9±5.1/<br>5.0±0.5 | 10.9±1.88/<br>11.69±2.06 | 25.69±11.02/<br>24.79±3.25 |
| Xiang 2014 (2) | China | N. D. | Aviscera Biosci. | 188/40 | 7.6 ± 1.2/<br>5.7 ± 0.5 | 7.8±0.89<br>/5.1±0.46 | 13.2 ± 7.1/<br>25.98 ± 2.88 | 25.6 ± 3.0/<br>23.8 ± 3.4 |
| Yildiz Kopuz 2022 (2) | Turkey | ADA (2022) | Phonix Pharm. | 38/38 | 9.4±3.3/<br>5.6±0.2 | 10.5±5.2<br>/4.9±0.4 | 2.57±0.44/<br>2.15±0.44 | 33.26 ± 5.23/<br>33.46 ± 6.58 |
| Zaki 2022 (2) | Egypt | N.D. | N.D. | 120/60 | 6.75±3.1/<br>3.91±1.01 | N.D. | 65.03±35.8/<br>40.3±4.34 | N.D. |
| Zamil 2019 (2) | Iraq | N.D. | MyBiosource | 80/80 | N.D. | 11.7±4.3<br>/5.1±0.7 | 3.9±3.2/<br>4.2±2.5 | N.D. |
| Zhang 2020 (2) | China | N. D. | Phoenix Pharm. | 350/194 | 8.7 ± 1.7/<br>5.0 ± 0.6 | N.D. | 8.03 ± 4.44/<br>7.91 ± 4.34 | 25.6 ± 3.73/<br>25.21 ± 3.29 |
| El Haddad 2019 (1) | Egypt | ADA (2014) | Wuhan Eiaab Science | 60/30 | 8.8 ± 1.0/<br>N. D. | 10.5±2.6<br>/4.7±0.4 | 0.18 ± 0.1/<br>0.16 ± 0.05 | 29.73 ± 2.89/<br>28.85 ± 1.66 |
| Li 2018 (1) | China | N.D. | Elabscience | 424/90 | 9.1 ± 1.9/<br>5.3 ± 0.59 | 9.2±3.2/<br>5.34±0.4 | 0.028±0.016<br>/0.056±0.034 | 25.9±3.35/<br>25.21±1.56 |
| Saadeldin 2018 (1) | Egypt | ADA (2015) | Glory Science | 61/8 | 9.6 ± 0.6/<br>5.5 ± 0.1 | 13.9±0.8<br>/5.4±0.1 | 0.04 ± 0.04/<br>0.47 ± 0.11 | 30.57 ± 1.29/<br>26.99 ± 1.6 |
| Yang 2020 (1) | China | WHO (1999) | Andy Gene Biotech. | 60/20 | 10.1±2.2/<br>5.6±0.32 | 9.4±2.7/<br>4.68±0.4 | 0.019±0.008<br>/0.037±0.02 | 24.9±2.58/<br>22.2±1.4 |
| Zhang 2014 (1) | China | N. D. | BioVendor | 64/30 | 9.4 ± 2.2/<br>5.4 ± 0.5 | 8.41±2.0<br>/4.81±0.58 | 0.019 ±<br>0.004/ 0.04 ± 0.003 | 22.61 ± 2.71/<br>22.45 ± 2.61 |

Noting the country where the study was conducted is relevant since it is known that T2DM differentially affects different populations (Hu 2011). Also noting the ELISA kit used to assess circulating levels of irisin is relevant, since different kits use different antibodies with varying degrees of specificity and affinity for irisin (Albrecht et al. 2015). Lastly noting the BMI is relevant not only because obesity is a primary risk factor for the development of T2DM (Nguyen et al. 2011), but also because diabetes is associated with loss of muscle mass and a more sedentary lifestyle. Indeed, an increase in BMI is expected to naturally lead to decreased irisin levels, thus constituting a potential confounding factor. Ideally, irisin levels should be compared between T2DM patients and healthy control participants within the same BMI class.

While the overall trend of the correlation between irisin levels and T2DM is similar across all included studies (Fig. 3), a significant amount of variability in several aspects of the reported data (Table 1) required stratification and sub-grouping to produce a reliable meta-analysis. Inconsistencies across studies were found mainly in: (1) average irisin concentrations, which ranged from a hundredth of a nanogram per milliliter (ng/mL) to thousands of ng/mL (Table 1); (2) the ELISA kits used to detect irisin; (3) statistical description and analysis of the data, for example, some studies reported the data as means +/− standard deviation or standard error, other studies reported the medians +/− IQR); (4) diagnostic criteria used to define the T2DM group – in some cases, the diagnostic criteria were not specified.

**Figure 3.**
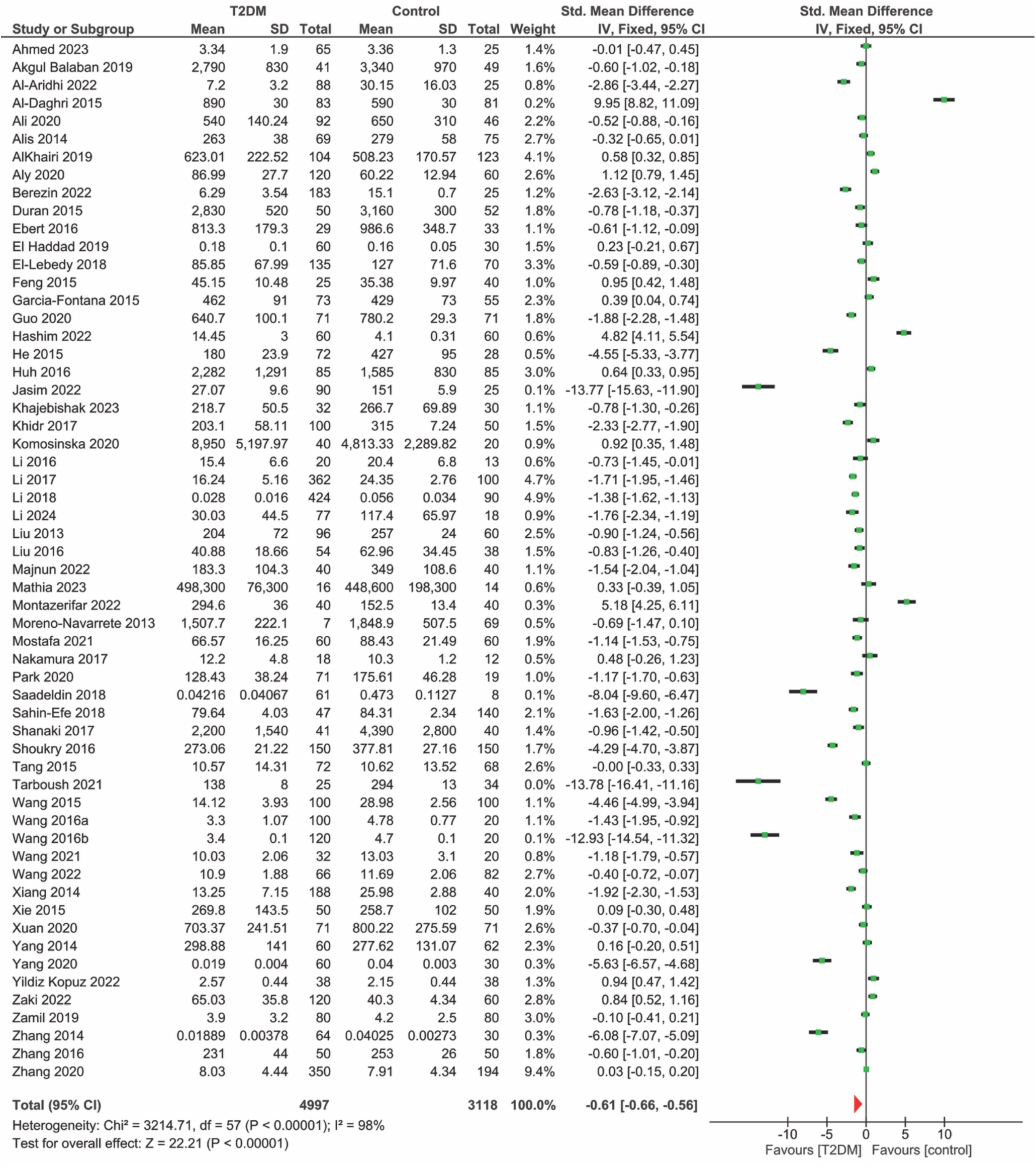
Forest plot of standardized mean differences in irisin concentration between diabetic (T2DM) and healthy control subjects across all studies included in this systematic review. 95% CI = [– 0.66 –0.56]. I2=98%. χ^2^ = 3214.71 (p <0.00001). Single-sample Z-test yields Z = 22.2 (p < 0.00001).

**Figure 4.**
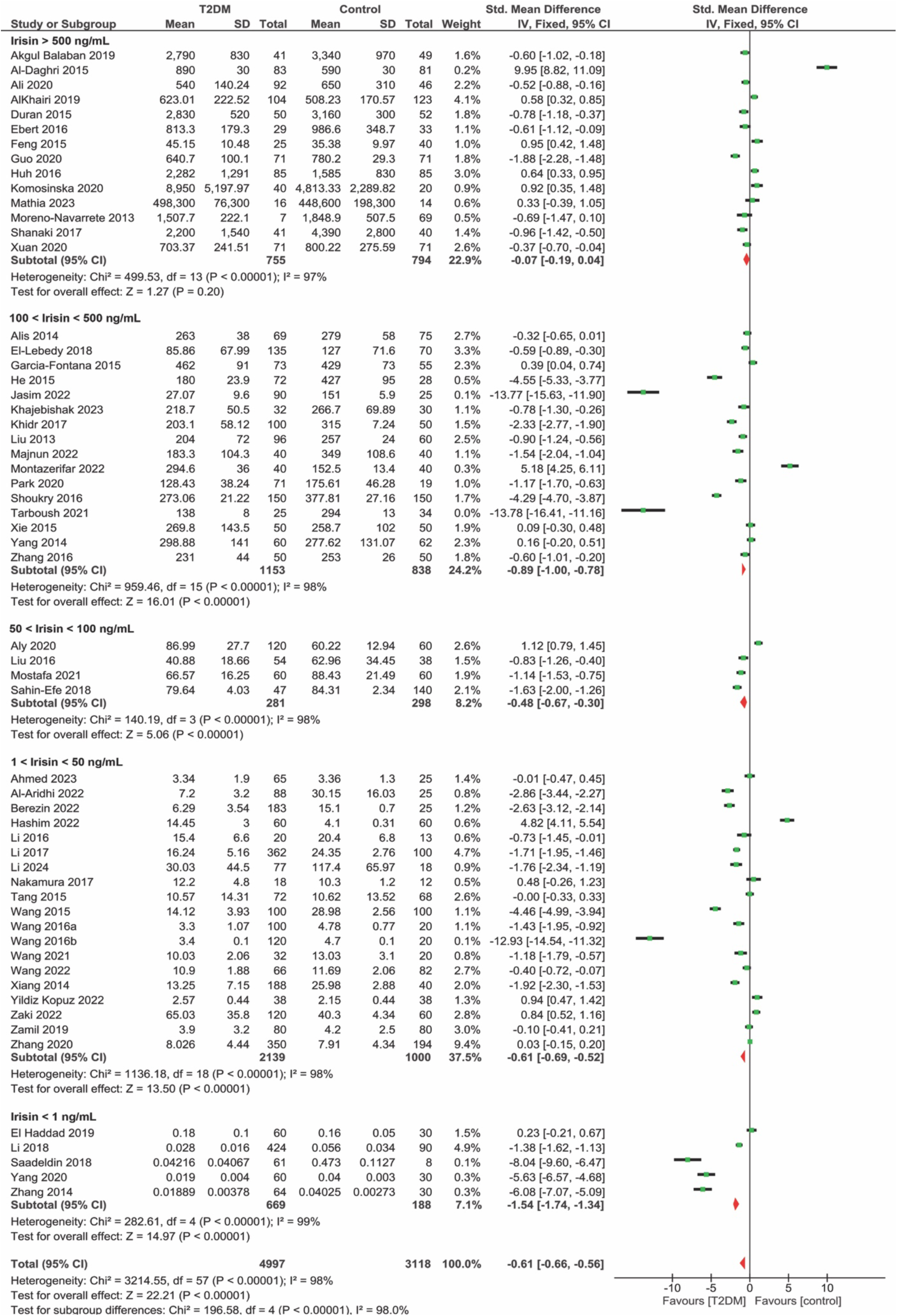
Forest plot of standardized mean differences between diabetic and healthy control subjects stratified by average absolute irisin concentration reported in the control group. Upon dividing all the studies in five subgroups (defined as: Subgroup 1: irisin <1 ng/mL, Subgroup 2: irisin = 1.01-50.00 ng/mL, Subgroup 3: irisin = 50.01-100 ng/mL, Subgroup 4: irisin = 100.01-500.00 ng/mL and, Subgroup 5: irisin≥ 500.01 ng/mL), no change in heterogeneity is noted in any of the subgroup analyzed (I^2^ ≤ 97% for all subgroups). In this subgroup analysis, effect trends remain negative and in favor of lower irisin in T2DM subjects, but effect size does change with irisin subgrouping. The greatest effect size is noted in Subgroup 1 (irisin <1 ng/mL), while the smallest effect size is noted in Subgroup 5 (irisin 500 < ng/mL). p < 0.00001 for all χ2 heterogeneity tests; p < 0.00001 for all one sample Z-tests except for the 500 < irisin concentration subgroup where p = 0.2.

### Irisin Levels Are Decreased in T2DM

When a meta-analysis is carried out across all included studies, the calculated 95% confidence interval (CI = –0.66, –0.56) for the standardized mean difference between irisin concentrations of T2DM subjects and control subjects appears to indicate that T2DM subjects have significantly lower irisin levels than those of healthy controls (Fig. 3). Eleven studies of the 58 included in the meta-analysis (Al Daghiri 2015, AlKhairi 2019, Aly 2020, Feng 2015, Garcia-Fontana 2015, Hashim 2022, Huh 2016, Komosinska 2020, Montazerifar 2022, Yildiz Kopuz 2022, and Zaki 2022) have an effect size 95% CI in favor of higher irisin concentrations in healthy controls compared to T2DM subjects. Similarly, when a Z-test was performed to calculate the significance of the effect size a p-value < 0.00001 was returned with a Z = 22.21 (Fig. 3).

Reported irisin concentrations varied from study to study by orders of magnitude. Indeed, as a result of the variability in reported irisin concentrations, *I^2^* – a measure of heterogeneity – is 98%, and the χ^2^ test for heterogeneity is significant as well (p < 0.00001). Although the effect size of T2DM on irisin concentration across all included studies is statistically significant, exceedingly high measures of heterogeneity indicate that these results need to be analyzed in subgroups to ensure calculated effect sizes are not dependent on variability in absolute irisin concentrations and other confounding variables (Fig. 3).

Subgroup analysis of absolute irisin concentrations does not aid in decreasing heterogeneity. For all subgroups, I^2^ ≥97% and the χ2 tests for heterogeneity yield p < 0.00001 for all subgroups (Fig. 4). Although heterogeneity is significantly affected by subgrouping, effect sizes are notably different across subgroups. The highest effect size is calculated in the <1 ng/mL irisin concentration subgroup (95% CI = [-1.74, –1.34]; Z = 14.97, p < 0.00001). The lowest effect size is noted in the subgroup 500 < ng/mL irisin (95% CI = [-0.19, 0.04]; Z = 0.61, p = 0.54). Of the 11 studies with 95% CI favoring healthy controls, five are located in the 500 < ng/mL irisin concentration subgroup (Al Daghiri 2015, AlKhairi 2019, Feng 2015, Komosinska 2020, and Mathia 2023) (Fig. 4).

### BMI is not a confounding factor

Nine studies present in the meta-analysis did not include BMI data for either healthy controls or T2DM subjects, so they were not included in this subgroup analysis. Once again, due to great variation in reported irisin concentrations, heterogeneity was exceedingly high (*I^2^ ≥* 97% for all subgroups). χ^2^ test for heterogeneity yields p < 0.00001 for all subgroups. The 95% confidence intervals for overall effect size only marginally depend on BMI classification when normal BMI participants are compared to obese (high BMI) participants (Fig. 5). Indeed, obese (–0.96 [-1.12, –0.8]) and normal (–0.97 [-1.05, –0.88]) BMI subgroups had approximately equal effect sizes and 95% confidence intervals; the effect of T2DM on irisin levels was significant in both subgroups since one-sample Z-tests for effect size indicate Z-test statistics are 21.82 and 11.70 for normal and obese BMI subgroups, respectively. In contrast, the subgroup of overweight participants displayed the lowest, though still significant, effect size (–0.33 [– 0.42, –0.25]) of T2DM on circulating irisin levels, suggesting that a slight increase in BMI may somehow interferes with the biological process(es) that link T2DM and circulating irisin levels (Fig. 5). Meta-regression for BMI was not conducted due to the extreme variability in reported irisin concentrations and the lack of raw BMI data.

**Figure 5.**
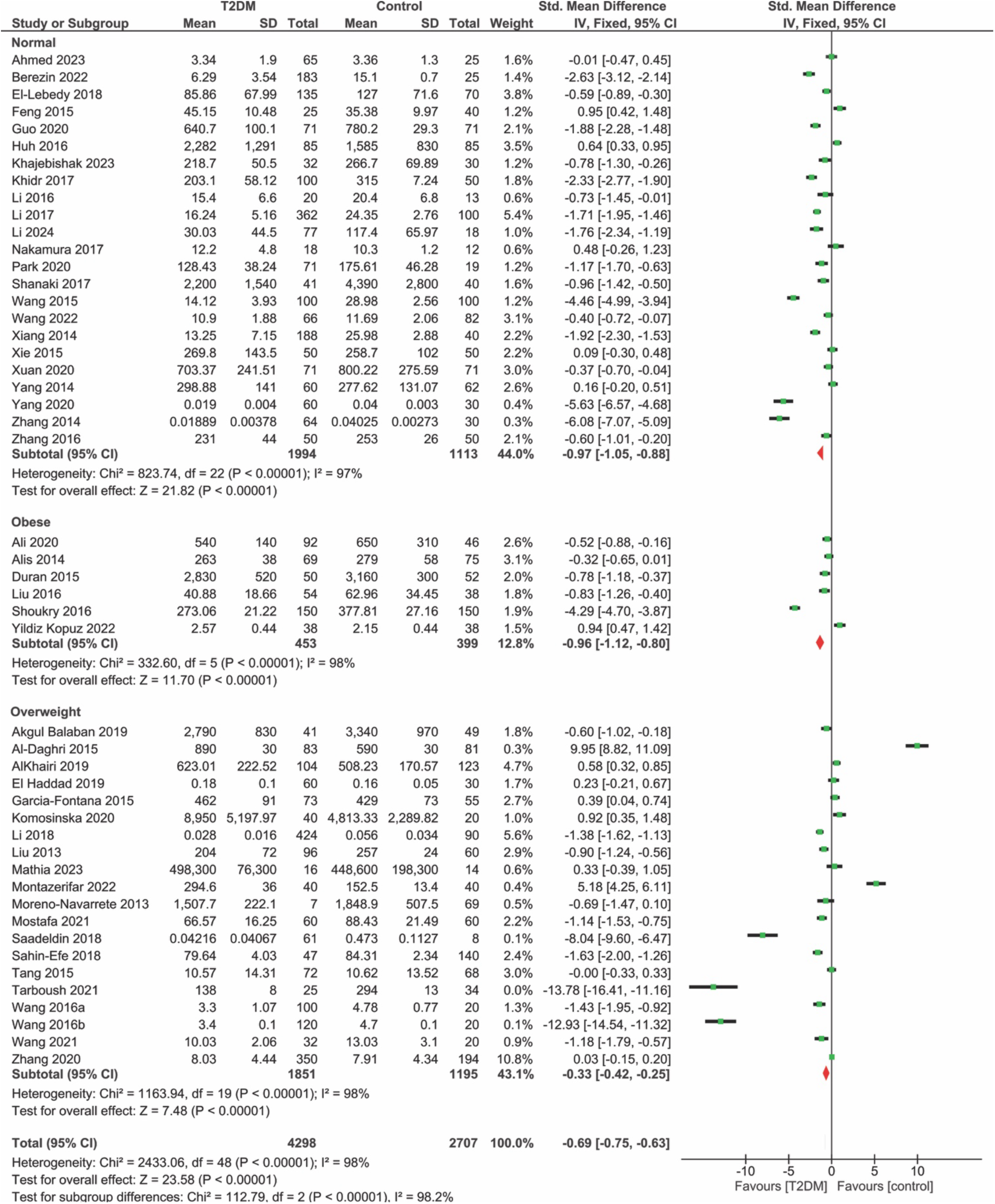
Forest plot of standardized mean difference between diabetic and healthy control subjects stratified by BMI classification of control subjects. The included studies were divided into three subgroups: Normal (BMI = 18.5 – 24.9), Overweight (BMI = 25-29.9) and Obese (BMI > 30). None of the studies contained either underweight (BMI < 18.5) or severely obese (BMI > 39) participants. Upon stratification based on BMI, heterogeneity remains high (I^2^ ≤ 97% for all subgroups). Effect trends remain negative and in favor of lower irisin in T2DM subjects, but effect size does change with BMI subgrouping. The greatest effect sizes are approximately equal among normal (–0.97 [-1.05, – 0.88]) and obese (–0.96 [–1.12, –0.8]) BMI subgroups. The lowest effect size is noted in the overweight subgroup (–0.33 [–0.42, –0.25]).

### Country of residence might be a confounding factor

Subgrouping by country yielded three subgroups: *China, Egypt*, and *Other*. All subgroups exhibited high heterogeneity in both the I2 and χ2 tests for heterogeneity (p < 0.00001 for χ2 tests; I2≥ 98%). In all subgroups, effect sizes and confidence intervals in their entirety favor lower irisin concentrations in T2DM subjects compared to healthy controls. However, differences in effect size are noted between country subgroups. The *China* group has the greatest calculated effect size and 95% CI (–0.91 [-0.99, – 0.84]), while the *Other* subgroup exhibited the lowest effect size and 95% CI (–0.20 [-0.29, –0.11]) (Fig. 6). One sample Z-tests for overall effect size yield significant p-values, less than 0.00001, for all country subgroups, yet the *China* subgroup has the greatest Z-test statistic at 22.96 compared to 9.22 and 4.36 for the *Egypt* and *Other* subgroups, respectively. This is likely due to the larger sample size of both studies and subjects in the *China* subgroup (Fig. 6).

**Figure 6.**
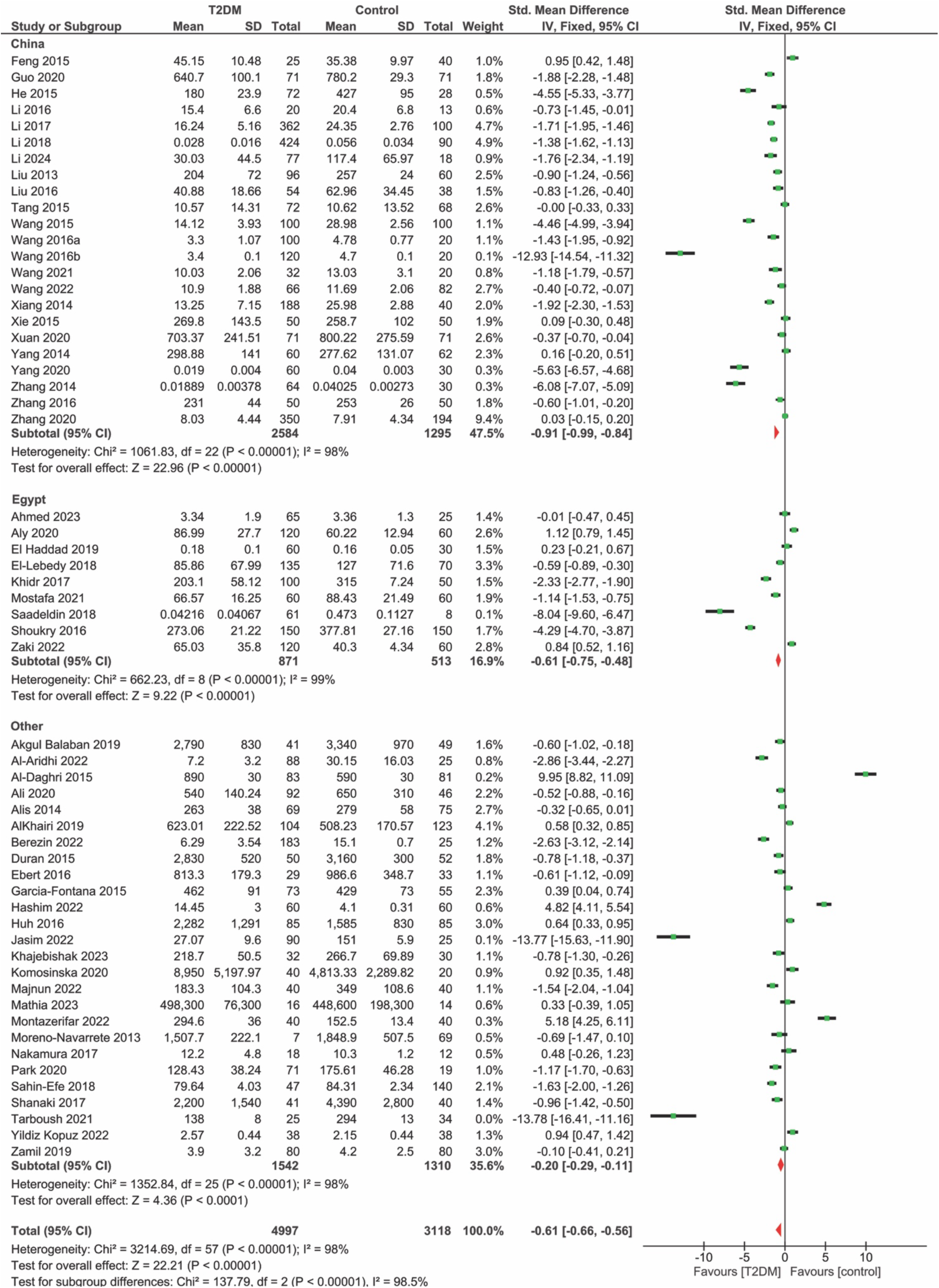
Forest plot of standardized mean differences between diabetic (T2DM) and healthy control subjects stratified by country. Studies were divided in three country subgroups: China, Egypt, and Other. Heterogeneity remains high in each subgroup (I^2^ ≥ 98% for all subgroups). Effect trends remain negative and in favor of lower irisin in T2DM subjects, but effect size does change with country subgrouping. The greatest effect size is calculated in the China subgroup and the lowest in the Other subgroup. p < 0.00001 for all χ2 heterogeneity tests and one-sample Z-tests for overall effect size.

## CONCLUSIONS

We have analyzed 58 original research articles published between 2011-2024 and carried out a meta-analysis to address the question: are circulating levels of irisin affected by T2DM? Meta-analysis of the available data shows a trend towards a decrease in irisin concentrations in T2DM subjects compared with healthy controls. Despite great variability across the studies examined, which decreased the statistical power of our meta-analysis, the finding that circulating irisin levels consistently trend to a reduction in T2DM patients compared with healthy controls, regardless of BMI, strongly suggest a direct relationship between T2DM and irisin that is not secondary to diabetic sarcopenia. In this regard, it should be noted that irisin not only is released by skeletal muscle during physical activity, but can also promote muscle hypertrophy (Reza et al., 2017), further stressing the importance of exercise for T2DM patients.

An important, technical issue with the literature systematically reviewed here, is that the average irisin concentrations across the included articles were greatly variable, thus resulting in statistical heterogeneity in the data set analyzed. This variability in reported irisin concentrations has been noted before by other researchers and attributed to the methods used to detect irisin, which rely on polyclonal antibodies of uncertain quality (Albrecht et al., 2015). ELISA is the primary method to detect irisin levels in plasma or serum samples. This meta-analysis includes studies that used ELISA kits from at least 12 different companies, many of which sell multiple ELISA kits with different detection ranges. Albrecht et al. discuss the possibility of aspecific cross-reacting proteins with polyclonal anti-irisin antibodies (Albrecht et al., 2015). However, their analysis relied on SDS-PAGE followed by western blotting rather than native PAGE. The denaturation of irisin by SDS-PAGE possibly alters its affinity for the polyclonal antibodies used in irisin detection ELISA kits, which detect the native protein, thus leaving the question open. In addition, it should be noted that of the five antibodies tested by Albrecht et al. (2015), four were problematic in that they were either discontinued (Aviscera/Phoenix; G-067-52), were not employed in commercially available ELISA kits (Cayman-14625, BioVision/BioCat-AP8746b-AB), or ELISA kits employing them were available but not tested in this study (Phoenix; G-067-16). Albeit with limitations, the study by Albrecht et al. reveals an important problem in the current methods used to evaluate irisin concentrations in biological fluids; at the moment, no consensus has been reached on the validity of ELISA for irisin quantification.

An alternative to ELISA for irisin quantification, as reported by Jedrychowski et al. (Jedrychowski et al., 2015) is mass spectrometry, which provides a highly effective and more precise quantification of irisin concentrations but may be more expensive for studies seeking to obtain high sample sizes. With mass spectrometry, Jedrychowski et al. conclude that irisin is indeed present in human plasma and that individuals undergoing aerobic exercise have elevated levels of irisin compared with sedentary individuals (Jedrychowski et al., 2015).

In conclusion, while a clear link between irisin and T2DM emerges from our study, the limitations mentioned above weakens the statistical significance of such finding. Nonetheless, the idea that decreased irisin levels in T2DM patients might be, at least in part, responsible for the observed pathology, is fascinating and deserves further investigation.

## CONFLICT OF INTEREST

The authors declare no conflict of interest.

## Notes

### Competing Interest Statement

The authors have declared no competing interest.

### Summary of Updates

Updated to include articles published in 2023-2024

